# Membrane Proteome of *Phytophthora parasitica* Zoospores: How Does Sensing Occur?

**DOI:** 10.1101/2025.03.28.645888

**Authors:** C.A. Lupatelli, A. Seassau, M. Magliano, M.L. Kuhn, A. Rey, M. Poët, L. Counillon, E. Evangelisti, P. Thomen, A. Attard, X. Noblin, E. Galiana

## Abstract

*Phytophthora* plant pathogens rely on motile biflagellated zoospores to efficiently locate and colonise host tissues. While rhizospheric signals guiding zoospore movement toward roots are known, the protein composition of membranes mediating these responses remains unclear. Here, we used liquid chromatography with tandem mass spectrometry (LC-MS/MS) and proteomic data mining to analyse membrane fractions from the flagella and cell bodies of *Phytophthora parasitica* zoospores. Major classes of membrane proteins (receptors, transporters and enzymes) were identified and their subcellular distribution between flagella and cell bodies quantified. Immunolocalization revealed that while most membrane proteins are evenly distributed, a subset localizes to the flagella, suggestive of specialized roles in sensing and movement regulation, particularly for sterol recruitment and ion flux variations. These findings advance our understanding of protein-mediated dispersal and host targeting by zoospores and support the hypothesis that zoospores use polarized signal perception mechanisms for environmental sensing and movement.

## 1 Introduction

Microbial species integrate a variety of environmental signals that shape their distribution and ultimately determine the microbiome composition within a given biotope. For pathogenic species, especially those in the oomycete class, responding to environmental factors is critical for exploring a microenvironment and establishing disease. Climatic forces, such as wind and water currents, are key to disseminating these pathogens, especially in soil and aquatic ecosystems^1^. Additionally, the chemical and physical properties of the biotope create conditions that can either facilitate or inhibit microbial exploration^2^. Within this complex environment, interactions between the microbiome and host plants play a crucial role, as host-emitted signals can stimulate chemotactic movement, trigger germination and sporulation, or initiate host penetration^2,3,4^.

In *Phytophthora* plant pathogens, the spread of disease in soil relies on motile zoospores that actively swim toward host roots^5,6^. Unicellular zoospores (6–10 µm diameter) propel themselves using two flagella (10–20 µm length) in a characteristic ‘run-and-tumble’ motion, where coordinated beats drive movement, and directional shifts involve a brief posterior flagellum pause^7^. Morphologically, zoospores are polarized cells, with two distinct flagella: an anterior decorated with mastigonemes, and a posterior with whiplash structure^7,8^.

Zoospores employ multiple taxis mechanisms to detect and integrate environmental cues critical for host localization and infection. Positive rheotaxis and negative geotaxis enable *Phytophthora* species to swim against currents and remain near the soil surface^3^. Navigation is further guided by electrotaxis, chemotaxis, and ion currents generated by plant roots. For instance, *P. sojae* and *P. cinnamoni* zoospores are attracted to plant exudates^9,10^. Potassium ion gradients, generated by soil activities, also influence zoospore swimming and aggregation^11,12^. In addition, internal signalling from pioneer zoospores plays a role in aggregating *P. parasitica* zoospores^13^.

Several molecular mechanisms that enable zoospores to detect and adjust their movement in response to external signals have been previously identified^3^. Potential chemoreceptors include G-protein-coupled receptors (GPCRs). For example, *P. infestans* and *P. sojae* mutants that lack the Gα subunit exhibit disrupted swimming patterns and lose chemotaxis toward glutamic acid and daidzein, respectively ^14,15^. Moreover, transformed *P. sojae* with altered leucine-rich repeat receptor-like kinases (RLKs) show impaired chemotaxis to daidzein, suggesting a role for RLKs as chemoreceptors^16^. The availability of *Phytophthora* genome sequences, coupled with advanced comparative omics approaches, raises the possibility for more systematic research into the perception and response of pathogen zoospores to signals in their environment.

In the present study, a proteomics approach was used to investigate the sensory repertoire of zoospores of *P. parasitica*, a polyphagous pathogen that infects a wide range of hosts^17^. We focused on three key questions: What is the plasma membrane (PM) protein repertoire that zoospores use to perceive and transduce external stimuli and navigate towards host plants? Do novel components within this repertoire reveal new insights into zoospore sensing? Are molecular sensors asymmetrically distributed between the flagella and the cell body, suggesting a specialization of roles?

To address these questions, we generated enriched membrane fractions from both the flagella and cell bodies of zoospores, to enable targeted analysis of these structures. Using liquid chromatography-tandem mass spectrometry (LC-MS/MS), the full zoospore membrane protein repertoire was qualitatively described and then the differences between the two fractions were quantified. Among the repertoire, we selected and characterized candidates such as enzymes, transporters, and receptors, investigated their subcellular localisation and used bioinformatics to investigate their putative roles in sensing and signal processing. Our findings shed light on the molecular basis of zoospore directed navigation and host signal detection.

## 2 Materials and Methods

### 2.1 Isolation of flagella and cell bodies from zoospore suspensions

The mycelium of *P. parasitica* (=*P. nicotianae*^17^) was cultured in a V8 liquid medium and subsequently incubated on 2% water agar for seven days to induce sporulation. Release of zoospores (10^6^ cells ml^-1^) was achieved in water^11^. Flagella (F) were separated from cell bodies (CB) by vortexing zoospore suspensions for two minutes. White light microscopy was used to check the transition of swimming zoospores to non-motile cysts and fluorescent microscopy with Tubulin Tracker Green staining was used to visualize flagella and cell bodies in suspension. The CB fractions were pelleted by centrifuging the deflagellated-cell suspension (3000 *g*, 5 min, RT). The supernatant was subjected to a second centrifugation (3000 g) to remove any CB contamination. The F fractions were pelleted at 31,000 *g* (30 min, 4°C).

### 2.2 Preparation of membrane-enriched fractions

We enriched four replicates of CB and F fractions in membrane (M) proteins using the Mem-PER Kit (ThermoFisher, Waltham, MA USA) and adapting the protocol for Suspension of Mammalian Cells. The CB and F pellet fractions were resuspended in 1 mL of the Cell Wash Solution buffer. After centrifugation at 1000*g* (3 min, RT) for CB fractions and 17000*g* (15 min, 4C°) for F fractions, the pellets were resuspended in the Permeabilization Buffer then the Solubilization Buffer. At each step F and CB pellets were resuspended in 200 µl and 500 µl of solution, respectively, incubated and gently mixed for 15 min at 4°C, and centrifuged at 17000*g* (15 min, 4°C). Supernatants, enriched in membrane fractions, were recovered in the Solubilization Buffer, and stored at −80°C. The protein concentration of the membrane-enriched fractions (M-F and M-CB) was determined by spectrophotometry (660 nm) using the Pierce™ BCA Protein Assay Kit with an ionic detergent compatibility reagent (ThermoFisher). The quality of fraction separation and membrane protein enrichment was confirmed by immunoblotting for the mastigoneme protein PPTG_02441, localized on the membrane of the anterior flagella.

### 2.3 Liquid chromatography and mass spectrometry analysis

The method is detailed in Supplementary Data 1. Briefly, four biological M-F and M-CB replicates were fractionated by electrophoresis through SDS-polyacrylamide (10%) gels, stained with Coomassie blue and sectioned into several bands (1 mm^3^). Each band was subjected to an acetonitrile washing, reduced with 10 mM DTT in 50 mM NH₄HCO₃ for 30 min at 56°C, and alkylated with 55 mM iodoacetamide in 50 mM NH_4_HCO_3_ for 20 min at 25°C in the dark. The proteinaceous content of bands was digested overnight at 37°C in a solution containing 25 mM NH_4_HCO_3_, 5 mM CaCl_2_, and 12.5 ng/µL sequencing-grade modified trypsin. The resulting peptides were extracted with acetonitrile. The samples were dried and desalted using OMIX C18 pipette tips (100 µL, A57003100) (Agilent, Santa Clara, CA, USA).

Samples were analysed using a nanoUHPLC system (nanoElute) coupled to a TimsTOFpro mass spectrometer (Bruker Daltonics, Germany). Each sample (10 µl) was separated on a reverse-phase C18 column with an integrated CaptiveSpray Emitter (75 µm ID x 250 mm, 1.6 µm, Aurora Series with CSI, ionOpticks, Australia). The TimsTOFpro mass spectrometer was operated with the CaptiveSpray nano-electrospray ion source. MS and MS/MS data were acquired in positive polarity using the Parallel Accumulation-Serial Fragmentation (PASEF) Data Dependent Acquisition (DDA) mode. Peptides were detected over a mass range of 100 to 1700 m/z, with a target intensity of 20,000 and an intensity threshold of 2500. The acquired DDA spectra were initially examined using Data Analysis software (version 5.3, Bruker Daltonics, Germany). Subsequently, the data were analysed against the proteome predicted from the *P. parasitica* 310 genome with PEAKS Studio (version Xpro, Bioinformatics Solutions)^18^. Only proteins identified with an FDR of 1%, with at least one unique peptide, were selected.

The relative abundances of the proteins in the M-F and M-CB replicates were measured using the PEAKS Q label-free quantification method. Only proteins with an FDR of less than 1% and a fold change greater than 2 were selected. To be considered for quantification, a minimum of one peptide was required, and peptides had to be identified in both groups and detected in at least two samples per group.

The proteomic data are available in the PRIDE database under the accession number PXD059087.

### 2.4 Data analysis: identification and quantification

The initial protein identification list generated with PEAKS (6516 proteins) was refined. Proteins were retained if they were identified by at least two MS/MS spectra across all four biological replicates, whether from the cell body or flagella fractions. The reduced protein list was further filtered using the TMHMM membrane-spanning helix predictor^19^, the NetGPI GPI-anchored proteins predictor^20^ and the SignalP 5.0 signal peptide residue predictor^21^. SignalP 5.0 predicted signal peptides in secreted and putative peripheral proteins associated with the membrane via protein-protein interactions. The refined list contained 1069 predicted zoospore membrane or secreted proteins. The protein families, names and Gene Ontology (GO) annotations were obtained from the UniProt database^22^. Classification of membrane proteins into three main categories (receptors, transporters and enzymes) was performed as described by Almén *et al.* (2009)^23^ and by finding functional annotations in the Transporter Classification DataBase (TCDB)^24^, the Enzyme classification (EC) system and other references in the literature^25^. Proteins lacking annotations were not further analysed. The remaining proteins with relevant biological functions were classified as ‘Other’.

### 2.5 *In silico* functional and structural annotation

Proteins of interest were functionally annotated with UniProt and SMART^26^ to identify the protein family and domain architecture. When available, predicted models of 3D structures were retrieved from AlphaFoldD or generated through AlphaFoldD2^27^. Phyre2 was used to identify and retrieve the best-aligning crystal structures of the proteins of interest^28^. At the same time, Chimera was used for structural alignment and calculation of the root-mean-square deviation (RMSD)^29^. To cluster members of a specific protein family, sequences were retrieved from GenBank using a three-step process: (i) initial homology searches, against the predicted proteome of *Phytophthora parasitica*, were conducted using BLASTP (threshold 1e-06) followed by (ii) searches against other public predicted proteomes from the REFSEQ_protein databank. (iii) Sequences were analysed with the CDvist tool^30^ to search for protein multi-domains, selecting sequences that displayed consistent patterns and orders of domain organization and exhibited at least 25% similarity to each other.

### 2.6 Immunolocalization experiments

Immunohistochemistry was performed with commercially produced Rabbit polyclonal antibodies targeting peptides for PPTG_02441 (CGNPGRLRTPEIAS), PPTG_09633 (CQDEYNDTIPTGGD), PPTG_07417 (CTGYGRNPTSWSLNDEN) and PPTG_13136 (CDLDNANTDGPLTEYLTKDA). Zoospores were first fixed using 1% (V/V) glutaraldehyde for 2 hours at 4°C, washed in phosphate-buffered saline (PBS) and treated with a blocking solution containing 3% bovine serum albumin in PBS, pH 7.2, for 30 min. They were then incubated for 3h at 4°C with each primary antibody, diluted 1:100, washed three times with PBS and incubated for 1h at RT with a FluoProbes 547H Donkey Anti-Rabbit IgG diluted 1:200 (Interbiotech – BioScience Innovations). Samples were rinsed three times and mounted in Fluoroshield (Sigma, St. Louis, MA USA). Image acquisition was performed using a Zeiss LSM 510 META confocal laser scanning microscope (Zeiss, Germany).

For immunoblotting, samples (2 µg) were separated in 4–20% precast polyacrylamide gels (Bio-Rad), transferred onto a nitrocellulose membrane and stained with Ponceau red. The membranes were incubated and washed sequentially in PBS in the presence of 5% (w/v) non-fat dry milk, a primary rabbit polyclonal antibody (1:2000), a goat peroxidase-conjugated IgG directed against rabbit immunoglobulins (1:5000), and a chemiluminescent substrate for peroxidase. The bound antibodies were detected using the FusionX apparatus (Vilber, Marne-la-Vallée, France) and the ECL Western Blotting Substrate protein labelling kit (Promega).

## 3 Results and Discussion

### 3.1 Production of cell body and flagella membrane protein-enriched fractions

To investigate the overall composition and distribution of zoospore membrane proteins, we purified cell body (CB) and flagellar (F) fractions from swimming *P. parasitica* zoospores (Fig. 1). The homogeneity of the purified fractions was assessed using light microscopy and tubulin staining (Fig. 1a-d). The F fraction comprised elongated hair-like and curved structures observed in detached flagella (Fig. 1c), while the CB fraction comprised round cells typical of encysted zoospores (Fig. 1d). We further processed the F and CB fractions to obtain the membrane protein-enriched fractions M-F and M-CB. As a quality control, we analysed M-F and M-CB by immunodetection using a polyclonal antibody raised against the mastigoneme protein PnMas1 (PPTG_02441), which localizes to the anterior flagellum of zoospores. The antibody decorated the anterior flagellum and partially the cell body at the flagellum base (Fig. 1e), likely corresponding to the diffusion barrier regulating protein migration toward the flagellar structure^31,32^. Western Blotting revealed a single band at 64 kDa corresponding to the predicted molecular weight of PnMas1 in samples corresponding to intact zoospores (used as a positive control) and the M-F fraction, but not in the M-CB fraction (Fig. 1f). Together, these results confirm the efficiency of our extraction method and the purity of the M-F and M-CB fractions.

**Figure 1.**
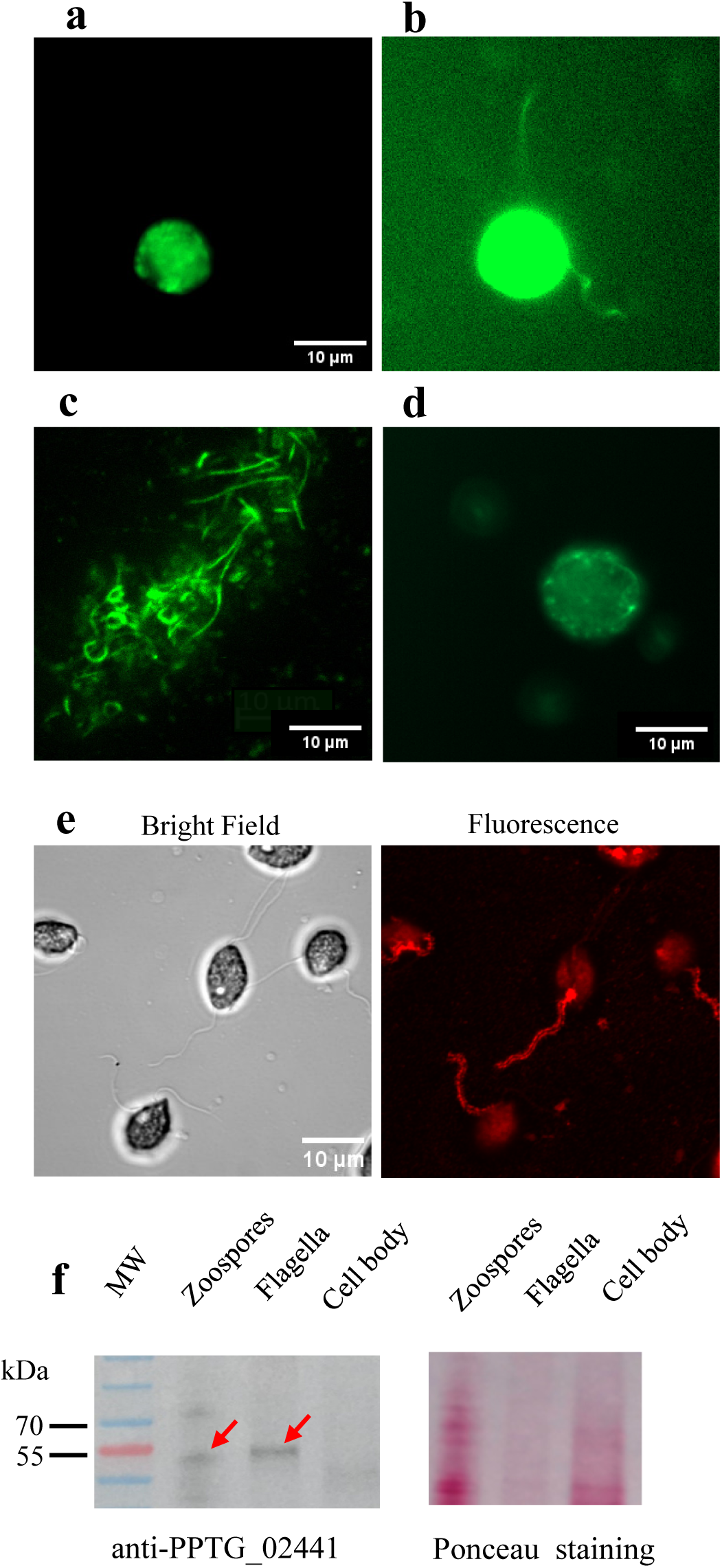
Cell body and flagella membrane-enriched fractions. (a, b) Tubulin Tracker Green labelling of an entire zoospore not overexposed and overexposed to enhance the visibility of flagella. (c, d) show separated flagella and cell body fractions. (e) Bright field (left) and fluorescence (right) views of zoospores labelled with antibodies against PPTG_02441. Observations were performed at confocal microscope using the 63x objective. (f) Immunoblot analysis using the anti-mastigoneme protein, PPTG_02441, detected at the expected calculated molecular weight (64kDa). Red arrows indicate the corresponding bands.

### 3.2 Membrane-associated proteins related to zoospore biology

LC-MS/MS analyses of the M-F and M-CB fractions identified 1,069 membrane-associated proteins (Supplementary Data 2). Of these, 843 proteins (70%) could be functionally classified as transporters, receptors, enzymes^33^ or ‘Other Classified Proteins’, which include proteins associated with specific aspects of *Phytophthora* biology (Fig. 2). The remaining 30% included proteins of unknown function and those unrelated to the defined categories, and were excluded from further analysis. We further mined the classified proteins for those potentially involved in zoospore sensing, signal transduction, and motion regulation.

**Figure 2.**
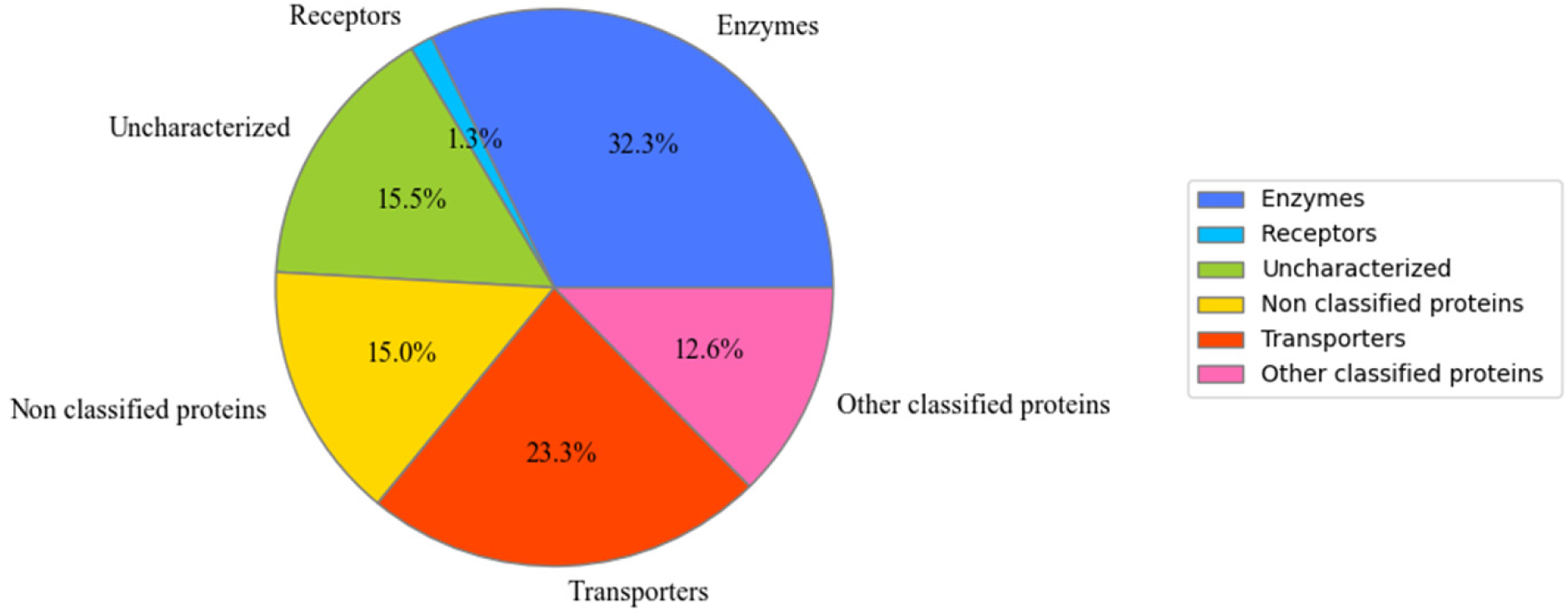
Categories of membrane proteins identified in zoospores. The main categories of membrane proteins detected were transporters, enzymes and receptors. Proteins with sufficient data but outside these categories were labelled as other classified proteins, while those with other unrelated functions were excluded. Notably, 15% of identified membrane proteins were unannotated and/or uncharacterized.

### Enzymes

Enzymes constituted the largest category, accounting for 32% of the classified proteins (Fig. 3a). Hydrolases made up more than 50% of identified enzymes, including 67 glycoside hydrolases (GH), with 9 cellulases and 60 peptidases. Classification of carbohydrate-active enzymes^34^ and *in silico* analysis of genes encoding cell wall-degrading enzymes in the *P. parasitica* genome^35^ revealed that the zoospore GH repertoire spans eleven hydrolase families (GH1, GH5, GH17, GH30, GH31, GH35, GH38, GH47, GH63, GH72, GH81), highlighting its capability to degrade various carbohydrates. Members of the GH1 family are thought to modify the *P. parasitica* cell wall rather than that of the host^35^. Members of the GH30 family (PPTG_08507, PPTG_08509) were annotated as glucosylceramidases and may play specialized roles in cellular recognition processes by modulating variations in the glucosylceramide head groups of their substrates, similar to functions observed in other eukaryotes^36^. Among the 60 peptidases identified, significant functional diversity was observed. Some were classified as metallopeptidases belonging to the thimet oligopeptidase family^37^ (e.g., PPTG_03858), which are known in *Trypanosoma brucei* for their role in the generation of quorum sensing signals^38^ and so are potentially involved in sensing.

**Figure 3.**
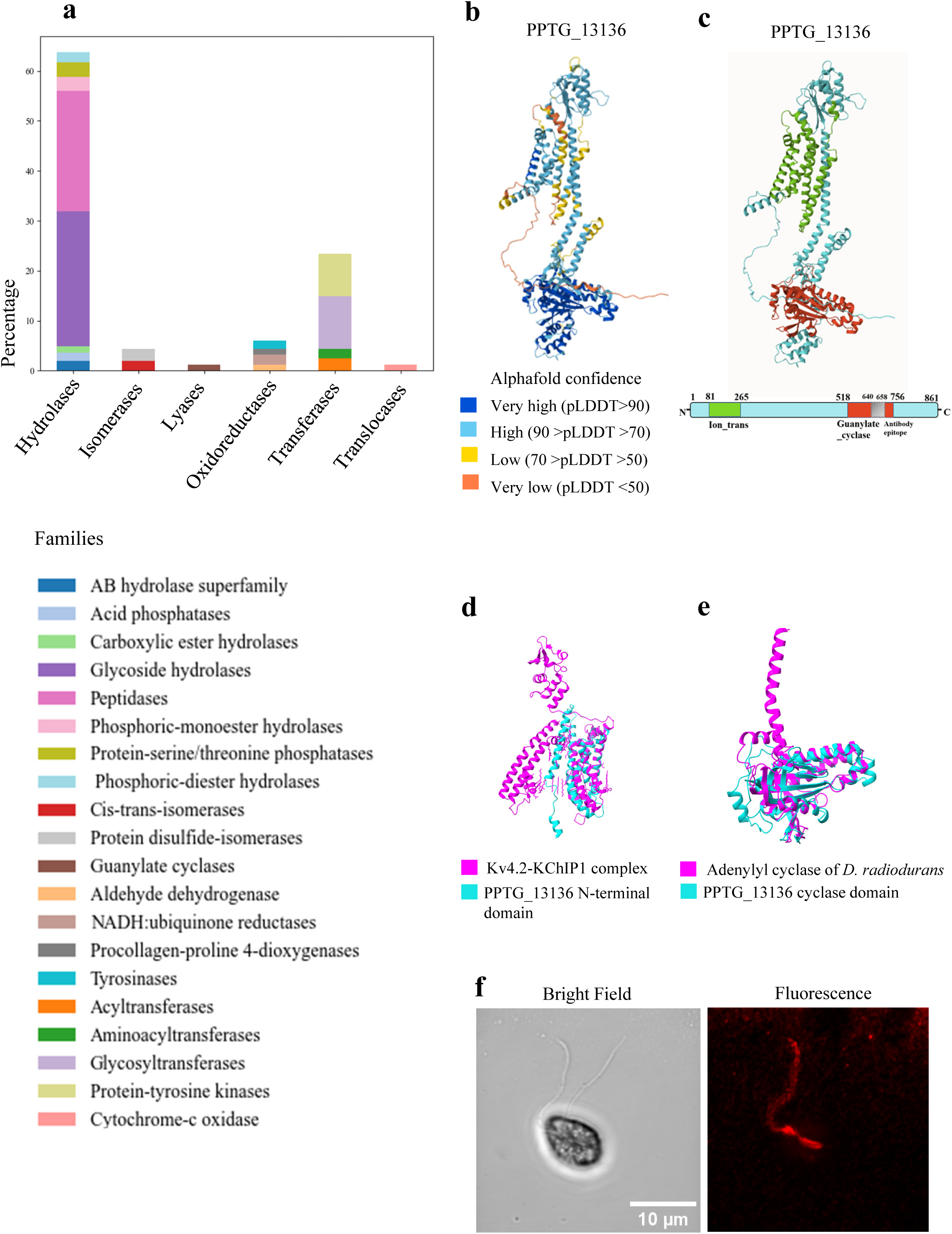
Zoospore membrane enzymes. (a) Distribution of enzyme families identified through membrane proteomic analysis of zoospores, presented as percentages. (b) AlphaFold model of PPTG_13136 with the pLDDT score indicating the accuracy of the prediction for each residue. (c) Domain architecture of PPTG_13136 indicating the position of the Ion_trans PFAM (green) and Guanylate cyclase domains (red). The same architecture was established for PPTG_13761. (d) Local alignment using ChimeraX of the 3D-structure prediction of the PPTG_13136 (cyan) N-terminal domain with a segment of the CryoEM structure of the human voltage-gated potassium channel, Kv4.2-KChIP1 complex (magenta, PDB: 2R9R). RMSD 12. (e) Superposition of the PPTG_13136 nucleotide cyclase domain (cyan) and Chain B of photoactivated adenylyl cyclase of *Deinococcus radiodurans* (magenta, PDB: 6FHT). RMSD 4. (f) Bright-field microscopy (left) and fluorescence confocal laser microscopy (right) images of zoospores stained for PPTG_13136. Images were obtained by using indirect immunofluorescence confocal laser microscopy.

Alongside hydrolases, 25% of the enzymes were identified as transferases, including 21 tyrosine kinases and 26 glycosyltransferases. The glycosyltransferase group, including β-1,3-D-glucan-(PPTG_08579, PPTG_13258, PPTG_13182) and cellulose-(PPTG_17902) synthases, may constitute preassembled enzymatic machinery that rapidly constructs the primary cyst wall post-deflagellation^39^. The 21 proteins annotated as tyrosine kinases exhibited characteristic sensing properties, with a typical organization reminiscent of the family of receptor-like kinases (RLKs): an N-terminal extracellular domain, a transmembrane domain, and a C-terminal kinase domain (KD)^40^. Notably, six of these (PPTG_00076, PPTG_01724, PPTG_01987, PPTG_09207, PPTG_14643, PPTG_16352) are distinguished by one to three Leucine Rich Repeat (LRR) domains in their extracellular region, which are crucial for protein binding and upstream signalling pathway activation^41^.

Finally, a small subset of lyases (2% of the enzymes) were identified as guanylate/adenylate cyclases (GCs/ACs), including PPTG_13136, PPTG_13761, and PPTG_08439, which are potentially crucial for zoospore sensing and signalling by catalysing cyclic GMP (cGMP) or cyclic AMP (cAMP) synthesis^42^. GCs and ACs are classified into soluble and membrane-bound types, with the former involved in signalling pathway functions like protein kinase activation and the latter categorized based on ligand specificity^43^. The proteins PPTG_13136 and PPTG_13761 annotated in the *P. parasitica* genome as voltage-gated ion channels, were found to be unique members of a bimodal membrane-bound protein family, presenting similarities with no other protein of *P. parasitica*. They share 38.75% sequence identity and each has two structurally independent domains. The N-terminal domain (PPTG_13136: residues 81-265; PPTG_13761: residues 81-226) includes an Ion_trans PFAM domain, characteristic of sodium, potassium, and calcium ion channels (Fig. 3b-c). These domains feature 5 and 3 transmembrane helices respectively, with segments containing typical motifs of the voltage sensor S4 segment (VSD) in voltage-dependent gated channels (Supplementary Fig.1a)^44^. Homology searches show similarity to human potassium voltage-gated channel subunits, and 3D structural predictions align these segments with the human shaker family voltage-dependent potassium channel and a subunit of the Kv4.2-KChIP1 complex (Fig. 3d). However, both proteins lack sequence homology in the regions corresponding to the pore helix, which is crucial for the gating mechanisms of K^+^ channels. The second domain encompasses residues 518-756 and 533-774 of PPTG_13136 and PPTG_13761 respectively, was annotated with a Guanylate_cyc PFAM domain (Fig. 3c and Supplementary Fig.1b). This domain, confirmed by Phyre2 modelling with high confidence, aligns closely with known 3D structures of adenylate and guanylate cyclases, suggesting functional similarity to guanylate/adenylate signalling cyclases (Fig. 3e). Taken together, the organization into two domains comprising a voltage sensor S4 segment (VSD) and a guanylyl cyclase (GC)/adenylyl cyclase (AC) suggests two potential functions for PPTG_13136 and PPTG_13761. They may act as ion channels with an enzymatic domain that plays a regulatory role, similar to certain plant cyclic nucleotide-binding domain (CNBD) superfamily ion channels, also found in ciliates and green algae^45^. Alternatively, they might act as ion current-regulated enzymes. This is akin to a novel adenylyl cyclase identified in *Paramecium* species, which features a VSD and a K^+^ channel pore localized at the ciliary level^46^, or the voltage-sensing phosphatases (VSPs) in choanoflagellates, where a VSD triggers phosphatase activity^47^. Here we identified similar bimodal proteins, exclusively in at least 90 other flagellated or ciliated organisms, within Stramenopiles and Alveolates (Supplementary Data 3 and Supplementary Fig.1c-d), suggesting that PPTG_13136 and PPTG_13761 might have a role also in motility-related processes. Supporting this hypothesis, immunolocalization revealed PPTG_13136 on *P. parasitica* zoospores, predominantly on a single flagellum, and possibly at its base (Fig. 3f). Further functional characterization is necessary to elucidate the role and regulatory mechanisms of these proteins.

### Transporters

Transporters were the second-largest heterogeneous category in the zoospore membrane proteome (23% of the total, n=249; Fig. 4a). Similar patterns have been previously observed in other flagellate species such as *Leishmania major*, *Trypanosoma brucei* and *Trichomonas vaginalis*, which have 247, 299 and 408 transporters, respectively^48^. This underscores the pivotal role of transporters in mediating the selective passage of molecules across the membrane of flagellated cells, thereby maintaining the electrochemical gradient essential for energy production and metabolic exchange, in particular in all aspects of the motility process, including environmental sensing, force generation and signalling^49^.

**Figure 4.**
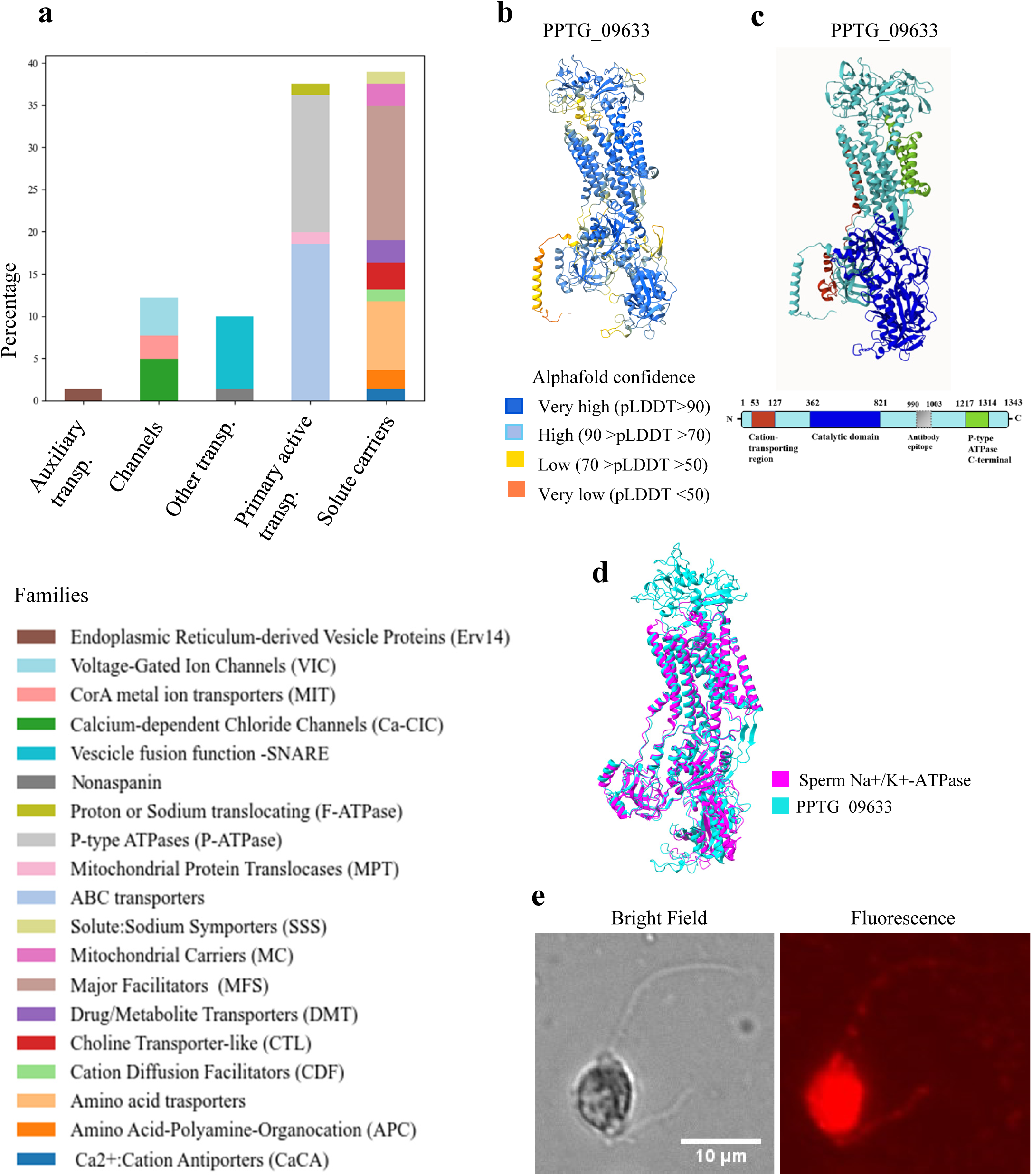
Zoospore membrane transporters. (a) Distribution of transporter families identified through membrane proteomic analysis of zoospores, presented as percentages. (b) AlphaFold model of PPTG_09633 with the pLDDT score indicating the accuracy of the prediction for each residue. (c) Schematic representation of PPTG_09633 domain architecture. The protein includes a cation-transporting P-type ATPase N-terminal region (53-127 aa) coloured in red, a catalytic domain (362-821 aa) in blue, and a P-type ATPase C-terminal region (1217-1314 aa) in green. (d) Tight structural alignment between the predicted 3D structure of PPTG_09633 (cyan) and the crystal structure of sperm Na^+^/K^+^-ATPase (magenta, PDB: 8ZYJ). RMSD 5. (e) Bright-field microscopy (left) and fluorescence confocal laser microscopy (right) images of zoospores stained for PPTG_09633.

Within this category, solute carriers comprised 38% of transporters, mainly from the major facilitator superfamily (n=35). Primary active transporters were similarly prevalent, representing 33% of the transporters, including P-type ATPases (n=36) and ABC transporters (n=41), along with a smaller subset of F-type ATPases (n=3). Among these, two subunits of the V-type proton ATPase were identified: the integral membrane subunit A (PPTG_05924) and the proteolipid subunit (PPTG_11905), which in zoospores localize to the spongiome membranes of the water expulsion vacuole (WEV), facilitating osmoregulation by powering water expulsion^50^.

Among the transporters, our analysis focused on the sodium-potassium ATPase, PPTG_09633, as it was the most abundant protein among all the 1069 identified membrane proteins (Fig. 4b). Eukaryotic Na^+^/K^+^-ATPases operate in the PM to maintain cellular homeostasis by transporting three Na^+^ ions out and two K^+^ ions into the cell, against their concentration gradients. These enzymes consist of an α-subunit with 10 transmembrane regions, which houses the catalytic and ion-binding sites, and a smaller β-subunit, which aids in membrane targeting and K^+^ ion access^51^. PPTG_09633 exhibited α-subunit characteristics (Fig. 4c) and *in silico* annotation associated it with K^+^ import and Na^+^ export functions across the PM. By screening the *P. parasitica* genome, two additional Na^+^/K^+^-ATPase α-subunits were identified, both of which were included in the list of membrane proteins. This finding aligns with estimates for *P. infestans*, which also presents two predicted Na^+^/K^+^-ATPase α-subunits^52^. No β-subunits were found, consistent with previous studies indicating their absence in fungi and most Protista^53^. The function of Na^+^/K^+^-ATPases in *Phytophthora* remains to be elucidated. Previous studies on *Pythium* species indicate how a putative P-type ATPase may pump K^+^ into the cell in low K^+^ environments^52^. In higher eukaryotic flagellate species, Na^+^/K^+^-ATPase functions have been widely characterized and are also associated with regulation of sperm flagellar beating^54,55,56^. Comparative alignment revealed high sequence conservation between PPTG_09633 and the mammalian sperm Na^+^/K^+^-ATPase isoform α4, with similar 3D-structural alignment, suggesting functional similarities (Fig. 4d). Moreover, PPTG_09633 was immunolocalized on both flagella and cell bodies (Fig.4e), showing that the Na^+^/K^+^-ATPase is distributed throughout the zoospore, contributing both to overall cell homeostasis and, on the flagella, potentially playing a role in motion regulation similar to the sperm Na^+^/K^+^-ATPase.

Finally, the results highlighted the critical function of pore-mediated membrane transport, demonstrated by the presence of diverse voltage-gated ion channels (n=10), comprising 14% of transporters and including a putative potassium channel (PPTG_12381). Voltage-gated channels are indispensable in flagellate behaviour, as demonstrated in *Chlamydomonas*, where specific voltage-gated calcium channels regulate photobehavioural responses^57^, and in human spermatozoa, where the CatSper2 voltage-gated calcium channel drives hyperactivation and vigorous flagellar movements, enabling sperm to penetrate the egg cell membrane during fertilization^58^. However, none of the *P. parasitica* proteins annotated as calcium channels (PPTG_16517, PPTG_13534, PPTG_11048) showed similarities to CatSper2.

### Receptors

Receptors represent only 1.3 % (n=14) of the identified membrane proteins (Fig. 5a).

**Figure 5.**
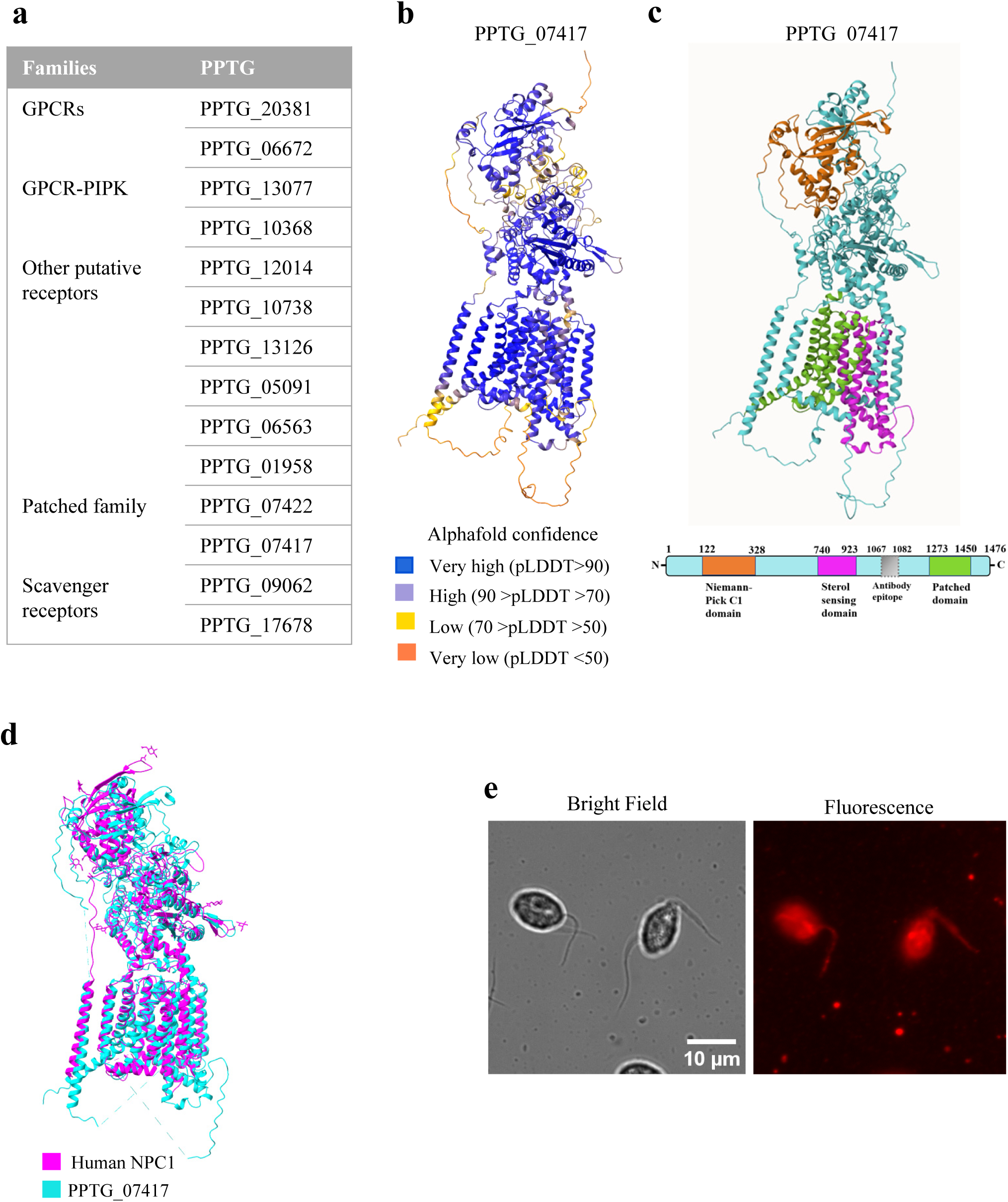
Zoospore membrane receptors. (a) Receptors identified through membrane proteomic analysis of zoospores. (b) AlphaFold model of PPTG_07417 with pLDDT score indicating the accuracy of prediction for each residue. (c) Domain structure of PPTG_07417, positioning the Niemann-Pick C1 domain (122-328 aa) coloured in orange, Sterol Sensing domain (740-923 aa) in magenta, and Patched domain (1273-1450 aa) in green. (d) Structural alignment of PPTG_07417 structure (cyan) with the crystal structure of full-length human NPC1 (magenta, PDB: 3JD8). RMSD = 14. (e) Bright-field microscopy (left) and fluorescence confocal laser microscopy (right) images of zoospores stained for PPTG_07417.

Interestingly we found only a few of the well-known components of major signalling pathways of microbial eukaryotic cells, such as G-protein-coupled receptors (GPCRs) and Patched family receptors from the Hedgehog (Hh) pathway^59^. While the human genome is estimated to encode approximately 950 GPCRs, of which 500 are odorant or taste receptors^60^, the genome of microorganisms such as *Dictyostelium* only encodes 55 GPCRs. Similarly, *Phytophthora* species (*P. infestans, P. sojae, P. ramorum*) harbour >50 GPCRs, 12 of which are fused to a phosphatidylinositol phosphate kinase (PIPK) domain^61^. One GPCR-PIPK proved to be involved in spore germination, hyphal elongation, sporangia cleavage and infection^62^. In our results, only two zoospore membrane proteins were identified as GPCRs (PPTG_20381 and PPTG_06672) and two as specific GPCR-PIPKs (PPTG_13077, PPTG_10638). This suggests that GPCRs may be less prevalent in the zoospore membrane so other sensory proteins likely contribute to the overall sensing capabilities.

Next, we identified two proteins, PPTG_07422 and PPTG_07417, that contain a sterol-sensing domain (SSD) and are involved in sterol sensing and uptake in *Phytophthora*^63^. SSD domains, approximately 180 amino acids in length, play a crucial role in the uptake of sterol substrates. These domains are found in proteins involved in sterol signal transduction, including sterol carrier proteins (SCP) like Niemann-Pick disease C1 (NPC1), essential for intestinal cholesterol absorption, and Patched (PTC) proteins^63,64^. *Phytophthora* species are unable to synthesize sterols and may use these receptors to get sterols from their environment and support their growth^64,69^. Four *P. capsici* genes, including the homologs of PPTG_07422 and PPTG_07417, which also encode SSD-containing proteins, have been found to play key roles, particularly in asexual reproduction and pathogenicity^63^. The identification of two proteins with SSDs on zoospore membranes raises the question of how zoospores contribute to *Phytophthora*’s detection and uptake of sterols, mechanisms that have only recently been explored^63^. Remarkably, PPTG_07417 the protein with a higher number of unique identified peptides, showed an N-terminal NPC1-like domain, then an SSD and a patched receptor domain (Fig. 5b-c). BLAST and 3D structural analyses revealed highly significant alignment between PPTG_07417 and NPC1 throughout their entire structures, especially within the SSD (Fig. 5d). These findings corroborate the study by Wang et al. (2023)^63^ suggesting a notable degree of evolutionary conservation of the NPC1 family. Immunolabelling of PPTG_07417 decorated one of the flagella and several cell body structures (Fig. 5e).

### Other classified proteins

Twelve percent of the proteins, categorized as Other classified proteins, contained sufficient information for functional classification based on UniProt and genomic annotations. Within this category, notable groups include proteins associated with lipid-related functions such as sensing, transport, signalling, rafts, and metabolism (n=30), *Phytophthora* pathogenicity (n=26), growth and signalling processes (n=19), protein-protein interactions (n=23), and a smaller number related to sperm functionality (n=3) (Fig. 6a).

**Figure 6.**
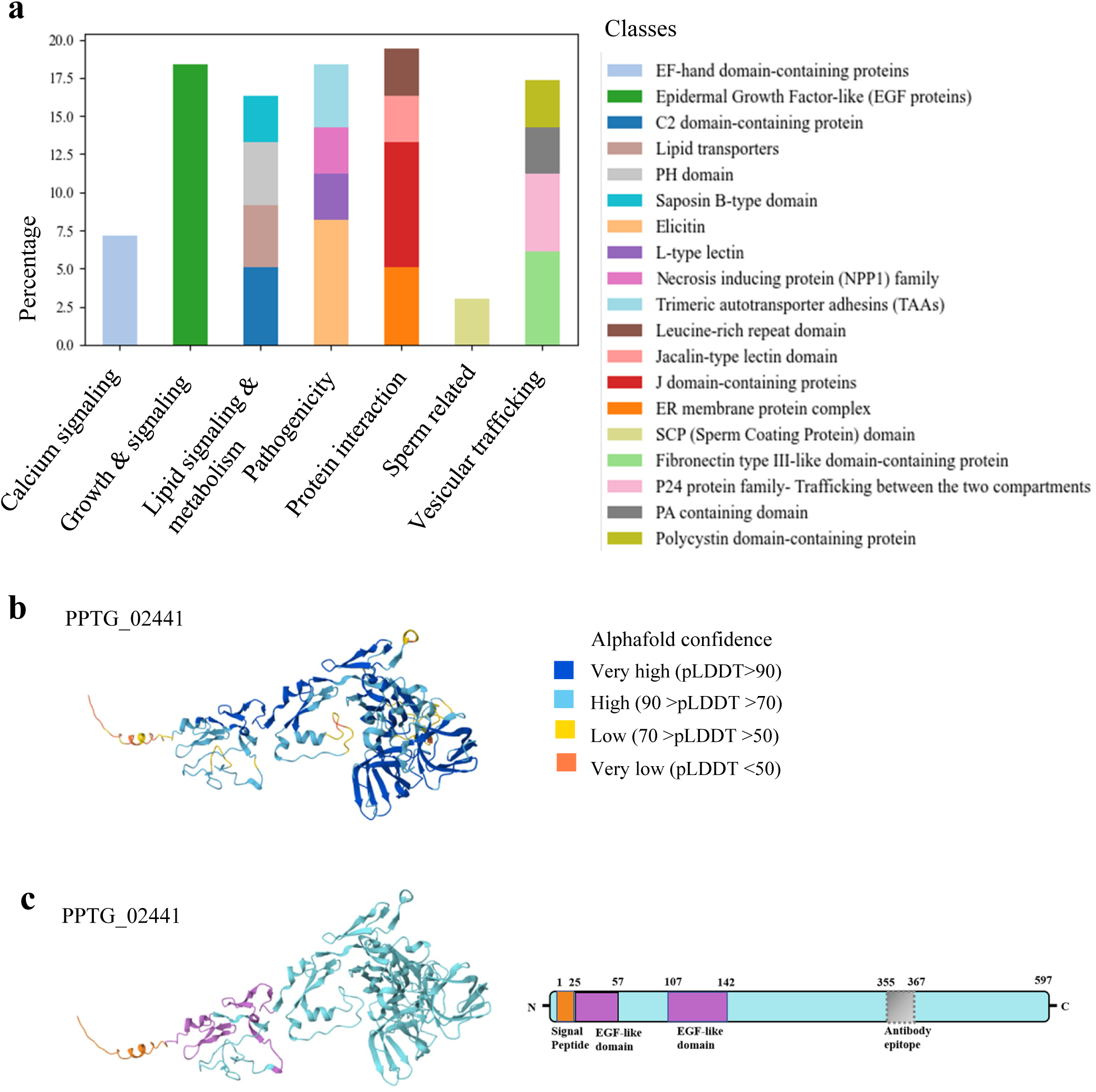
Annotation of other zoospore membrane proteins. (a) Distribution of membrane-related proteins of zoospores presented as percentages. (b) AlphaFold model of PPTG_02441 with pLDDT score indicating accuracy of prediction for each residue. (c) Domain structure of the mastigoneme protein PPTG_02441, showing at least 2 EGF-like domains (in violet) and a signal peptide (orange).

The functional cluster of 30 proteins related to lipid sensing, transport, and signalling is consistent with the identification of proteins containing the SSD, mentioned above. These findings highlight that lipid uptake and regulation are crucial functions in *Phytophthora* zoospores. Among this group, proteins featured domains such as START (PPTG_06151, PPTG_16174) and PH (PPTG_16268, PPTG_14048, PPTG_12937, PPTG_12938), each known for distinct contributions to lipid-related activities in animals. The START domain specializes in the transport of lipids and sterol binding and is vital for the shuttling of these molecules across cellular compartments^65^. The PH domain acts in membrane-associated signalling pathways by binding to phosphoinositides, essential components in signal transduction mechanisms that involve lipids^66^.

The group of proteins implicated in oomycete pathogenicity (n=26) includes several putative elicitins and NPP1 proteins. Elicitins, conserved extracellular proteins in *Phytophthora* and *Pythium*, function as microbe-associated molecular patterns (MAMPs) to trigger defence responses in various plant species^67^. Interestingly, some elicitins are anchored to the oomycete PM rather than being fully secreted^67^. In particular, this may be the case for PPTG_17779, PPTG_07488, PPTG_13458, PPTG_08951 and PPTG_12308, which are annotated as elicitins and exhibit a GPI-anchor site. Analysis of the NPP1 proteins (here PPTG_15231, PPTG_08017 and PPTG_07664), which are implicated in causing necrosis in plant leaves and roots and are noted for being secreted, reveals that these proteins primarily feature a signal peptide, as previously documented^68^. While the presence of these secreted proteins in membrane extracts could be due to contamination, we may also speculate that they are localized to the membrane’s external face as peripheral proteins bound via protein-protein interactions.

Within the category of other classified proteins, another subgroup (n=18) comprises proteins involved in growth and signalling, mostly characterized by cysteine-rich EGF-like domains. This group included the three known mastigoneme proteins^8^ (PPTG_07238, PPTG_02441, PPTG_03648) that exhibit a signal peptide and three to four EGF-like domains, and no transmembrane domains. These proteins have been speculated to participate in mechanosensation and motion regulation in flagellated species within the Stramenopile taxon^8,69^. Figure 6 (b-c) presents the 3D structure and the domain composition of PPTG_02441.

Among the 23 proteins involved in protein-protein interactions, we identified four putative leucine-rich repeat (LRR) domain-containing proteins (PPTG_07054, PPTG_07055, PPTG_00288, PPTG_12191). Although their specific functions are unclear, LRR domains are typically found in proteins involved in protein-protein interactions, including tyrosine kinase receptors and cell-adhesion molecules^16^.

Finally, three proteins (PPTG_08737, PPTG_00570, PPTG_08378) were annotated as harbouring a sperm coating domain (SCP, or CAP) and a signal peptide. CAP superfamily proteins are ubiquitous across all kingdoms of life, predominantly appearing in secreted forms, and play diverse roles in immune defence in plants and mammals, sperm maturation and fertilization, and fungal virulence^70^. Within the sperm context, certain CAP protein subfamilies modulate potassium channel activity, contributing to their functions in sperm maturation and motility^71,72^. The *P.parasitica* proteins may similarly influence flagellate zoospores, affecting ion transport mechanisms and ultimately motility.

### 3.3 Flagella versus cell body variation

A quantitative analysis was performed to compare M-CB and M-F fractions and showed that 785 proteins were differentially enriched between the cell body and flagella (Fig. 7a) (Supplementary Data 2). Of these 785 proteins, 710 proteins were more abundant in cell body fractions (ratio <0.5), while only 75 proteins were more abundant in the flagella (ratio >2). Using the same filtering criteria as for the whole repertoire (GPI, signal peptide, and transmembrane predictions), we identified 137 membrane-associated proteins in the cell body fraction, and 45 membrane-associated proteins in the flagella fraction (which corresponds to more than half of the proteins more abundant in the flagella). Unsurprisingly, M-CB exhibited a broader diversity of membrane proteins than M-F, including proteins extracted from the membranes of internal organelles that are absent in M-F.

**Figure 7.**
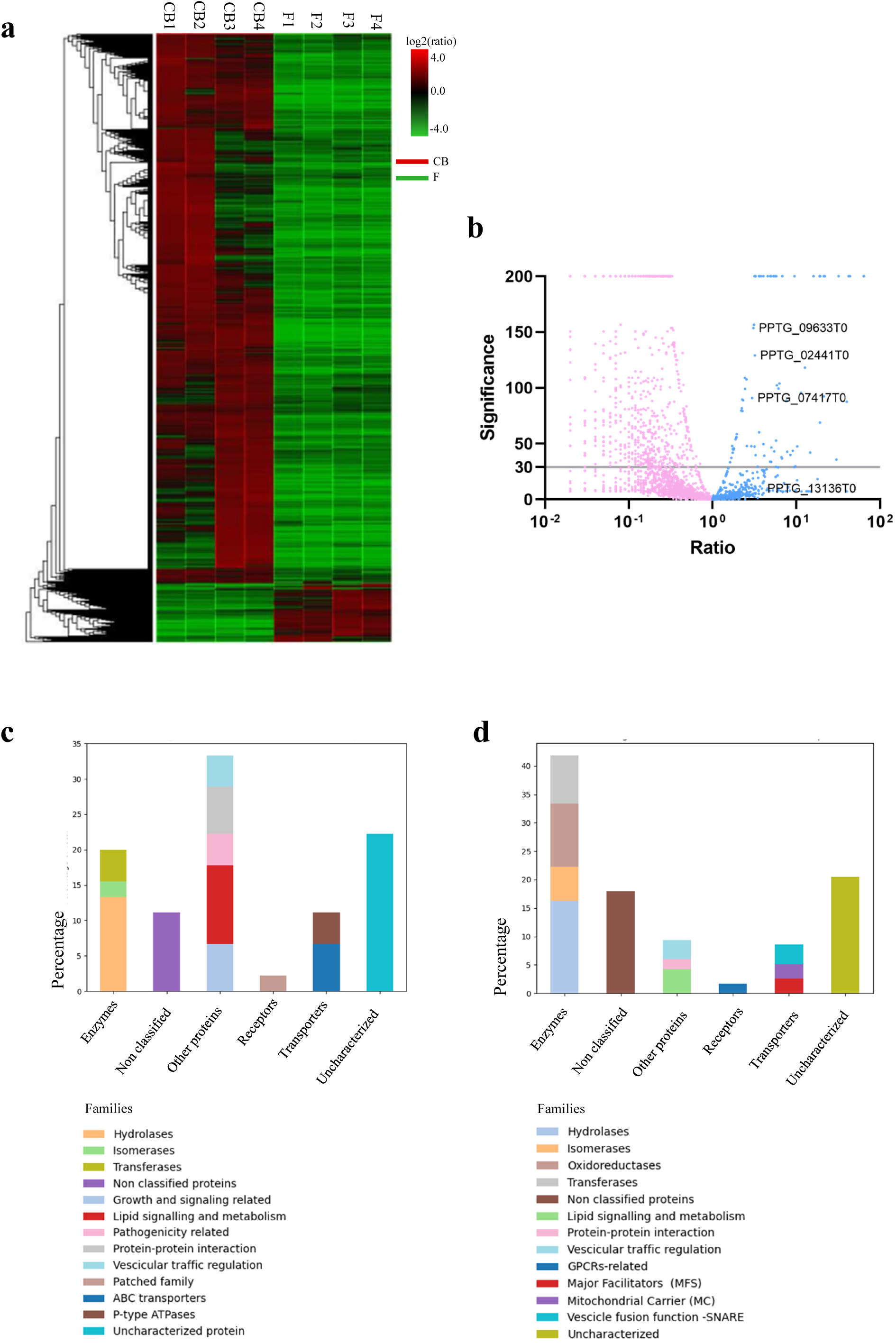
Distribution of proteins with significant differences in abundance between the four replicates of cell body and flagella samples. (a) Heatmap showing the proteins more abundant in the cell body (C1.1 to C4.1) or flagella replicates (F1.1 to F4.1). (b) Volcano plot of the different quantified proteins highlighting the proteins of interest that are more abundant in the flagella replicates. PPTG_13136 was not differentially distributed according to the quantification method adopted. (c) Membrane protein families more abundant in the flagella fraction. (d) Membrane protein families more abundant in the cell body fraction.

Compared to M-F (Fig. 7c), M-CB was particularly enriched with a variety of enzymes mainly involved in trafficking and metabolic processes, with only a minority of proteins involved in cell sensing and motion regulation (Fig. 7d). As expected, M-F was enriched with the three structural mastigoneme proteins, which we considered a quality criterion for the quantitative analysis. Lipid signalling and transporter proteins were also notably enriched in M-F, particularly the sterol-sensing domain-containing protein PPTG_07417, with a fold change of 2.97. Another protein, annotated as an ABC transporter domain-containing protein (PPTG_08986), has been identified through domain analysis as belonging to the ABC A-subfamily transporter, which is associated with lipid trafficking^73,74^. Two putative lipases, PPTG_01586 and PPTG_11537, were found to be more abundant in M-F. Collectively, these observations suggest there may be a specific mechanism for sterol/lipid sensing and uptake, regulated at the flagellar level, that warrants further investigation. Concerning the pathogenic process, PPTG_07664 was the sole highly enriched secreted protein effector identified in M-F^75^. Finally, the most represented protein among the 1069 identified proteins, namely the Na^+^/K^+^-ATPase PPTG_09633, was also more abundant in the M-F fraction (Fig. 7b).

A subset (30 %) of the proteins enriched in the M-F fraction were not predicted to be membrane-associated. Among these, we identified proteins with recognized roles in signalling chemotaxis in *Phytophthora*, including a G-protein α-subunit (PPTG_13453) and β-subunit (PPTG_18378)^14,62^. Interestingly, we also found the protein PPTG_11388, which harbours a VWA domain that mediates adhesion. This copine-like protein is very similar to CPNA in *Dyctostelium*, which is known to be involved in adhesion and chemotaxis^76^. Finally, we also found flagellar structural proteins such as α− and β-tubulins, and centriole and basal body proteins^77^. This is consistent with the observation that pieces of axonemes and axonemal protein remain associated with the flagellar membrane after non-ionic detergent extraction, reflecting sites of tight association between the axoneme and the flagellar membrane^32^.

While the quantification results highlight some minor differences between M-CB and M-F, the apparent differences may have been reduced because: (i) the relative abundance of flagellar proteins may have been underestimated due to their synthesis in the cell body and transient accumulation at diffusion barriers before transport to the flagella (as seen with the mastigoneme protein PPTG_02441); and (ii) proteins missing a peptide area value in one replicate were excluded from differential quantification, and this included proteins such as PPTG_13136 and PPTG_13761 that appeared more abundant in M-F and were confirmed by immunohistochemistry to be mostly localized on the flagella. Nevertheless, these results indicate a possible specialized role for the flagellar membrane in signal transduction and environmental responsiveness, emphasizing its functional adaptation beyond just motion. This aligns with findings in other flagellates like *Chlamydomonas*, where the flagellar membrane, though continuous with the cell’s PM, exhibits distinct protein variations linked to sensory functions^78^.

## 4 Conclusions

The study posed three primary questions aimed at uncovering the protein repertoire in the PM used by zoospores to perceive and respond to external stimuli, and to drive their motion toward host plants:

I. What is the overall PM protein repertoire used by zoospores during signal perception and motion adaptation? Our results revealed new insights into how *Phytophthora* zoospores may use their PM protein repertoire for sensing and motility, particularly for sterol recruitment and ion flux variations.
II. Are there novel components within the repertoire that reveal new mechanisms of zoospore sensing? PPTG_13136 and PPTG_13761 emerged as potential members of a new ciliary protein family, conserved across 90 flagellated or ciliated organisms. These proteins may play a role in regulating flagellar motion and the presence of a conserved voltage sensor domain suggests their potential involvement in ion-sensing processes. Although further investigation is needed to confirm their precise roles in zoospore sensing and motility, immunolocalization on *P. parasitica* zoospores revealed PPTG_13136 was predominantly localized on a single flagellum, strongly implying potential involvement in flagellar motility regulation.
III. To what extent are molecular sensors heterogeneously distributed in a zoospore? Our analysis identified an uneven distribution of molecular sensors, including the sterol-sensing protein PPTG_07417 and the S4 voltage domain-containing protein PPTG_13136. This suggests a possible polarization of function in zoospores that use flagella as sensory as well as motile organelles. This polarization likely plays a key role in regulating ion and lipid homeostasis, and sensing, transport, and signalling processes. These findings offer valuable insights into the role of flagella in zoospore motility and directional behaviour.

## 5 Acknowledgments

This research was funded by the French government through the Investissements d’Avenir program, managed by the French National Research Agency under the UCAJEDI project (grant number ANR-15-IDEX-01). We acknowledge support for Carlotta A. Lupatelli from a PhD grant provided by the Graduate Schools LIFE and SPECTRUM, and Academy 4 at Université Côte d’Azur. Additional funding was received from the French National Research Agency (grant number ANR22-CE20-0021) and from the Plant2Pro® Carnot Institute in the framework of its 2023 call for projects, supported by the ANR (agreement no. 20 CARN 0024). We also thank the PlantBIOs platform at Institut Sophia Agrobiotech for access to proteomic analysis equipment, funded through the SABLES platforms project with support from the European Union and the European Regional Development Fund. The authors thank Catherine Buchanan from the Office of International Scientific Visibility of Nice-Côte d’Azur University for editing the manuscript for grammar and English language expression.

## 6 Authors Contributions

EG, AA, XN, and CL conceived the study. EG, CL, AS, MM, and MK designed the experiments. CL, EG, MK, MM, AS, and AR performed the primary experiments. CL, EG, MM, and EE conducted data mining and bioinformatics. LC, MP, XN and EE contributed to the data interpretation and manuscript writing. EG, CL, and AA led the manuscript preparation. All authors reviewed, revised, and approved the final version of the manuscript.

---

**Supplementary Figure 1 Sequence properties of PPTG_13136 and PPTG_13761** (a) Alignment using ClustalW multiple alignment in BioEdit, of a part of the N-terminal Ion_trans PFAM domain of PPTG_13136 and PPTG_13761, compared with two human potassium voltage-gated channels, subfamily A member 4 (P22459) and member 3 (P22001). Shaded amino acids indicate AA homology; yellow boxes below sequences indicate the position of the TMD domains and of the matching with the helical segment S4, the voltage sensor of the voltage-dependent gated channel. (b) Alignment of the nucleotide cyclase domain of PPTG_13136 and PPTG_13761 with two bacterial photoactivated adenylyl cyclases (K9TLZ5 and Q9HZ23). (c) Table showing the number of proteins with domain organizations similar to PPTG_13136 and PPTG_13761 across different phyla. The highest BLAST scores are predominantly found in flagellate organisms, suggesting a role for these proteins in motility. The variability in protein numbers likely reflects the functional and structural complexity of each group, with ciliates exhibiting higher numbers (6-16), possibly due to the greater demands of their multiple ciliary structures. (d) Analysis of conserved domains in PPTG_13136. The proteins share with other similar bimodal proteins different conserved domains, corresponding here (from left to right) to the third transmembrane helix (4.8e-773 e-value), the S4 voltage sensor (4.3e-874 e-value), the fourth transmembrane helix (1.1e-1099 e-value) and part of the guanylate cyclase domain (4.5e-2375 e-value). Analysis was conducted using the MEME Suite (MEME Suite, Bailey et al., 2015).

## References

1. Thompson, S. E., Levin, S. & Rodriguez-Iturbe, I. Rainfall and temperatures changes have confounding impacts on Phytophthora cinnamomi occurrence risk in the southwestern USA under climate change scenarios. Glob. Change Biol. 20, 1299–1312 (2014).

2. Bassani, I., Larousse, M., Tran, Q. D., Attard, A. & Galiana, E. Phytophthora zoospores: From perception of environmental signals to inoculum formation on the host-root surface. Comput. Struct. Biotechnol. J. 18, 3766–3773 (2020).

3. Kasteel, M., Ketelaar, T. & Govers, F. Fatal attraction: How Phytophthora zoospores find their host. Semin. Cell Dev. Biol. 148–149, 13–21 (2023).

4. Situ, J. et al. Signal and regulatory mechanisms involved in spore development of Phytophthora and Peronophythora. Front. Microbiol. 13, 984672 (2022).

5. Walker, C. & West, P. van. Zoospore development in the oomycetes. Fungal Biol. Rev. 21, 10–18 (2007).

6. Mitchell, H. J., Kovac, K. A. & Hardham, A. R. Characterisation of *Phytophthora nicotianae* zoospore and cyst membrane proteins. Mycol. Res. 106, 1211–1223 (2002).

7. Tran, Q. D. et al. Coordination of two opposite flagella allows high-speed swimming and active turning of individual zoospores. eLife 11, e71227 (2022).

8. Hee, W. Y., Blackman, L. M. & Hardham, A. R. Characterisation of Stramenopile-specific mastigoneme proteins in Phytophthora parasitica. Protoplasma 256, 521–535 (2019).

9. Hua, C., Yang, X. & Wang, Y. *Phytophthora sojae* and soybean isoflavones, a model to study zoospore chemotaxis. Physiol. Mol. Plant Pathol. 92, 161–165 (2015).

10. Botha, T. & Kotzé, J. M. Exudates of avocado rootstocks and their possible role in resistance to Phytophthora cinnamomi.

11. Galiana, E., Cohen, C., Thomen, P., Etienne, C. & Noblin, X. Guidance of zoospores by potassium gradient sensing mediates aggregation. J. R. Soc. Interface 16, 20190367 (2019).

12. Appiah, A. A., van West, P., Osborne, M. C. & Gow, N. A. R. Potassium homeostasis influences the locomotion and encystment of zoospores of plant pathogenic oomycetes. Fungal Genet. Biol. 42, 213–223 (2005).

13. Galiana, E., Fourré, S. & Engler, G. Phytophthora parasitica biofilm formation: installation and organization of microcolonies on the surface of a host plant. Environ. Microbiol. 10, 2164–2171 (2008).

14. Latijnhouwers, M., Ligterink, W., Vleeshouwers, V. G. A. A., van West, P. & Govers, F. A Galpha subunit controls zoospore motility and virulence in the potato late blight pathogen Phytophthora infestans. Mol. Microbiol. 51, 925–936 (2004).

15. Hua, C. et al. A Phytophthora sojae G-Protein α Subunit Is Involved in Chemotaxis to Soybean Isoflavones. Eukaryot. Cell 7, 2133–2140 (2008).

16. Si, J., et al. *Phytophthora sojae* leucine-rich repeat receptor-like kinases: diverse and essential roles in development and pathogenicity. iScience 24, 102725 (2021).

17. Kamoun, S. et al. The Top 10 oomycete pathogens in molecular plant pathology. Mol. Plant Pathol. 16, 413–434 (2015).

18. Ma, B. et al. PEAKS: powerful software for peptide de novo sequencing by tandem mass spectrometry. Rapid Commun. Mass Spectrom. RCM 17, 2337–2342 (2003).

19. Krogh, A., Larsson, B., von Heijne, G. & Sonnhammer, E. L. L. Predicting transmembrane protein topology with a hidden markov model: application to complete genomes1. J. Mol. Biol. 305, 567–580 (2001).

20. Gíslason, M. H., Nielsen, H., Almagro Armenteros, J. J. & Johansen, A. R. Prediction of GPI-anchored proteins with pointer neural networks. Curr. Res. Biotechnol. 3, 6–13 (2021).

21. Almagro Armenteros, J. J., et al. SignalP 5.0 improves signal peptide predictions using deep neural networks. Nat. Biotechnol. 37, 420–423 (2019).

22. The UniProt Consortium. UniProt: the Universal Protein Knowledgebase in 2023. Nucleic Acids Res. 51, D523–D531 (2023).

23. Almén, M. S., Nordström, K. J., Fredriksson, R. & Schiöth, H. B. Mapping the human membrane proteome: a majority of the human membrane proteins can be classified according to function and evolutionary origin. BMC Biol. 7, 50 (2009).

24. Saier, M. H. et al. The Transporter Classification Database (TCDB): 2021 update. Nucleic Acids Res. 49, D461–D467 (2021).

25. McDonald, A. G. & Tipton, K. F. Enzyme nomenclature and classification: the state of the art. FEBS J. 290, 2214–2231 (2023).

26. Letunic, I. & Bork, P. 20 years of the SMART protein domain annotation resource. Nucleic Acids Res. 46, D493–D496 (2018).

27. Jumper, J. et al. Highly accurate protein structure prediction with AlphaFold. Nature 596, 583–589 (2021).

28. Kelley, L. A., Mezulis, S., Yates, C. M., Wass, M. N. & Sternberg, M. J. E. The Phyre2 web portal for protein modeling, prediction and analysis. Nat. Protoc. 10, 845–858 (2015).

29. Pettersen, E. F. et al. UCSF Chimera—A visualization system for exploratory research and analysis. J. Comput. Chem. 25, 1605–1612 (2004).

30. Adebali, O., Ortega, D. R. & Zhulin, I. B. CDvist: a webserver for identification and visualization of conserved domains in protein sequences. Bioinformatics 31, 1475–1477 (2015).

31. Hu, Q. & Nelson, W. J. The ciliary diffusion barrier: the gatekeeper for the primary cilium compartment. Cytoskelet. Hoboken NJ 68, 313–324 (2011).

32. Bloodgood, R. A. The Chlamydomonas Flagellar Membrane and Its Dynamic Properties. in 309–368 (Elsevier, 2009). doi:10.1016/B978-0-12-370873-1.00048-4.

33. Almén, M. S., Nordström, K. J., Fredriksson, R. & Schiöth, H. B. Mapping the human membrane proteome: a majority of the human membrane proteins can be classified according to function and evolutionary origin. BMC Biol. 7, 50 (2009).

34. Drula, E. et al. The carbohydrate-active enzyme database: functions and literature. Nucleic Acids Res. 50, D571–D577 (2022).

35. Blackman, L. M., Cullerne, D. P. & Hardham, A. R. Bioinformatic characterisation of genes encoding cell wall degrading enzymes in the Phytophthora parasitica genome. BMC Genomics 15, 785 (2014).

36. Dwek, R. A. et al. The fate and function of glycosphingolipid glucosylceramide. Philos. Trans. R. Soc. Lond. B. Biol. Sci. 358, 869–873 (2003).

37. Schoina, C. et al. Mining oomycete proteomes for metalloproteases leads to identification of candidate virulence factors in Phytophthora infestans. Mol. Plant Pathol. 22, 551–563 (2021).

38. Tettey, M. D., Rojas, F. & Matthews, K. R. Extracellular release of two peptidases dominates generation of the trypanosome quorum-sensing signal. Nat. Commun. 13, 3322 (2022).

39. Wang, M. C. & Bartnicki-Garcia, S. Synthesis of noncellulosic cell-wall β-glucan by cell-free extracts from zoospores and cysts of*Phytophthora palmivora*. Exp. Mycol. 6, 125–135 (1982).

40. Dievart, A., Gottin, C., Périn, C., Ranwez, V. & Chantret, N. Origin and Diversity of Plant Receptor-Like Kinases. Annu. Rev. Plant Biol. 71, 131–156 (2020).

41. Ng, A. & Xavier, R. J. Leucine-rich repeat (LRR) proteins. Autophagy 7, 1082–1084 (2011).

42. Karan, S., Frederick, J. M. & Baehr, W. Novel Functions of Photoreceptor Guanylate Cyclases Revealed by Targeted Deletion. Mol. Cell. Biochem. 334, 141–155 (2010).

43. Bassler, J., Schultz, J. E. & Lupas, A. N. Adenylate cyclases: Receivers, transducers, and generators of signals. Cell. Signal. 46, 135–144 (2018).

44. Noda, M. et al. Primary structure of Electrophorus electricus sodium channel deduced from cDNA sequence. Nature 312, 121–127 (1984).

45. Jegla, T., Busey, G. & Assmann, S. M. Evolution and Structural Characteristics of Plant Voltage-Gated K+ Channels. Plant Cell 30, 2898–2909 (2018).

46. Weber, J. H. et al. Adenylyl cyclases from *Plasmodium*, *Paramecium* and *Tetrahymena* are novel ion channel/enzyme fusion proteins. Cell. Signal. 16, 115–125 (2004).

47. Arcos-Hernández, C. & Nishigaki, T. Ion currents through the voltage sensor domain of distinct families of proteins. J. Biol. Phys. 49, 393–413 (2023).

48. Dean, P., Major, P., Nakjang, S., Hirt, R. P. & Embley, T. M. Transport proteins of parasitic protists and their role in nutrient salvage. Front. Plant Sci. 5, 153 (2014).

49. Jimenez, V. & Mesones, S. Down the membrane hole: Ion channels in protozoan parasites. PLoS Pathog. 18, e1011004 (2022).

50. Mitchell, H. J. & Hardham, A. R. Characterisation of the water expulsion vacuole inPhytophthora nicotianae zoospores. Protoplasma 206, 118–130 (1999).

51. Clausen, M. V., Hilbers, F. & Poulsen, H. The Structure and Function of the Na,K-ATPase Isoforms in Health and Disease. Front. Physiol. 8, 371 (2017).

52. Barrero-Gil, J., Garciadeblás, B. & Benito, B. Sodium, potassium-atpases in algae and oomycetes. J. Bioenerg. Biomembr. 37, 269–278 (2005).

53. Shahnazari, M., Zakipour, Z., Razi, H., Moghadam, A. & Alemzadeh, A. Bioinformatics approaches for classification and investigation of the evolution of the Na/K-ATPase alpha-subunit. BMC Ecol. Evol. 22, 122 (2022).

54. Jimenez, T., McDermott, J. P., Sánchez, G. & Blanco, G. Na,K-ATPase alpha4 isoform is essential for sperm fertility. Proc. Natl. Acad. Sci. U. S. A. 108, 644–649 (2011).

55. Tiwari, S. et al. Testis-Specific Isoform of Na+-K+ ATPase and Regulation of Bull Fertility. Int. J. Mol. Sci. 23, 7936 (2022).

56. Trötschel, C. et al. Absolute proteomic quantification reveals design principles of sperm flagellar chemosensation. EMBO J. 39, e102723 (2020).

57. Sanyal, S. K. et al. Characterization of Chlamydomonas voltage-gated calcium channel and its interaction with photoreceptor support VGCC modulated photobehavioral response in the green alga. Int. J. Biol. Macromol. 245, 125492 (2023).

58. Quill, T. A., Ren, D., Clapham, D. E. & Garbers, D. L. A voltage-gated ion channel expressed specifically in spermatozoa. Proc. Natl. Acad. Sci. U. S. A. 98, 12527–12531 (2001).

59. Nechipurenko, I. V. The Enigmatic Role of Lipids in Cilia Signaling. Front. Cell Dev. Biol. 8, 777 (2020).

60. Mykytyn, K. & Askwith, C. G-Protein-Coupled Receptor Signaling in Cilia. Cold Spring Harb. Perspect. Biol. 9, a028183 (2017).

61. Bakthavatsalam, D., Meijer, H. J. G., Noegel, A. A. & Govers, F. Novel phosphatidylinositol phosphate kinases with a G-protein coupled receptor signature are shared by Dictyostelium and Phytophthora. Trends Microbiol. 14, 378–382 (2006).

62. Hua, C. et al. GK4, a G-protein-coupled receptor with a phosphatidylinositol phosphate kinase domain in Phytophthora infestans, is involved in sporangia development and virulence. Mol. Microbiol. 88, 352–370 (2013).

63. Wang, W. et al. Sterol-Sensing Domain (SSD)-Containing Proteins in Sterol Auxotrophic Phytophthora capsici Mediate Sterol Signaling and Play a Role in Asexual Reproduction and Pathogenicity. Microbiol. Spectr. 11, e03797–22.

64. Wang, W., Liu, X. & Govers, F. The mysterious route of sterols in oomycetes. PLoS Pathog. 17, e1009591 (2021).

65. Tsujishita, Y. & Hurley, J. H. Structure and lipid transport mechanism of a StAR-related domain. Nat. Struct. Biol. 7, 408–414 (2000).

66. Lemmon, M. A. Pleckstrin homology (PH) domains and phosphoinositides. Biochem. Soc. Symp. 81–93 (2007) doi:10.1042/BSS0740081.

67. Derevnina, L. et al. Emerging oomycete threats to plants and animals. Philos. Trans. R. Soc. Lond. B. Biol. Sci. 371, 20150459 (2016).

68. Martins, I. M., Meirinho, S., Costa, R., Cravador, A. & Choupina, A. Cloning, characterization, in vitro and in planta expression of a necrosis-inducing Phytophthora protein 1 gene npp1 from Phytophthora cinnamomi. Mol. Biol. Rep. 46, 6453–6462 (2019).

69. Huang, J. et al. Structure-guided discovery of protein and glycan components in native mastigonemes. Cell 187, 1733–1744.e12 (2024).

70. Schneiter, R. & Di Pietro, A. The CAP protein superfamily: function in sterol export and fungal virulence. Biomol. Concepts 4, 519–525 (2013).

71. Guo, M. et al. Crystal structure of the cysteine-rich secretory protein stecrisp reveals that the cysteine-rich domain has a K+ channel inhibitor-like fold. J. Biol. Chem. 280, 12405–12412 (2005).

72. Gibbs, G. M. et al. Cysteine-rich secretory protein 4 is an inhibitor of transient receptor potential M8 with a role in establishing sperm function. Proc. Natl. Acad. Sci. U. S. A. 108, 7034–7039 (2011).

73. Albrecht, C. & Viturro, E. The ABCA subfamily--gene and protein structures, functions and associated hereditary diseases. Pflugers Arch. 453, 581–589 (2007).

74. Vasiliou, V., Vasiliou, K. & Nebert, D. W. Human ATP-binding cassette (ABC) transporter family. Hum. Genomics 3, 281–290 (2009).

75. Gijzen, M. & Nürnberger, T. Nep1-like proteins from plant pathogens: recruitment and diversification of the NPP1 domain across taxa. Phytochemistry 67, 1800–1807 (2006).

76. Buccilli, M. J. et al. Copine A Interacts with Actin Filaments and Plays a Role in Chemotaxis and Adhesion. Cells 8, 758 (2019).

77. Chen, X., Shi, Z., Yang, F., Zhou, T. & Xie, S. Deciphering cilia and ciliopathies using proteomic approaches. FEBS J. 290, 2590–2603 (2023).

78. Bloodgood, R. A. Sensory reception is an attribute of both primary cilia and motile cilia. J. Cell Sci. 123, 505–509 (2010).

