## Supplementary Fig.1 for "Membrane Proteome of *Phytophthora parasitica* Zoospores: How Does Sensing Occur?"

**a**

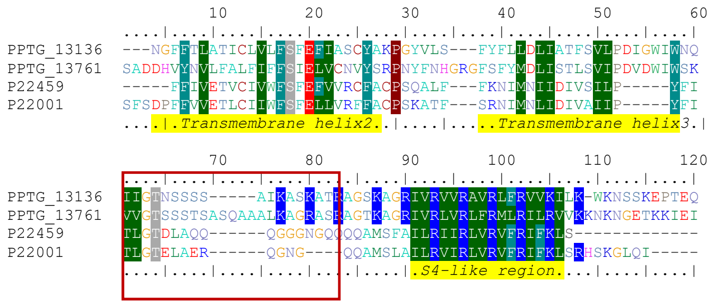


**b**

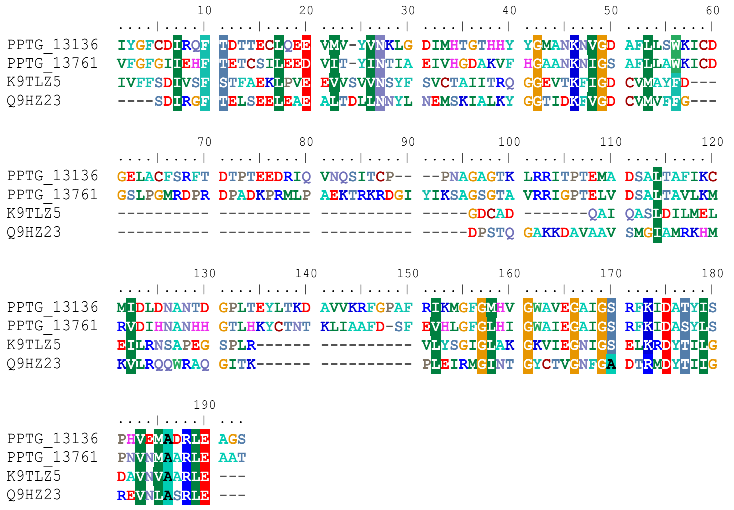


**c**

| Phylum/Super group | Class/Clade | Protein number |
|---|---|---|
| Stramenopiles | Oomycetes | 2 to 4 |
| | Pelagophytes | 2 |
| | Diatoms | 1 |
| Alveolata | Apicomplexans | 1 to 2 |
| | Ciliates | 6 to 16 |
| | Dinoflagellates | 3 |
| Chlorophyta | Green algae | 1 |
| Haptista | Haptophytes | 1 |

**d**

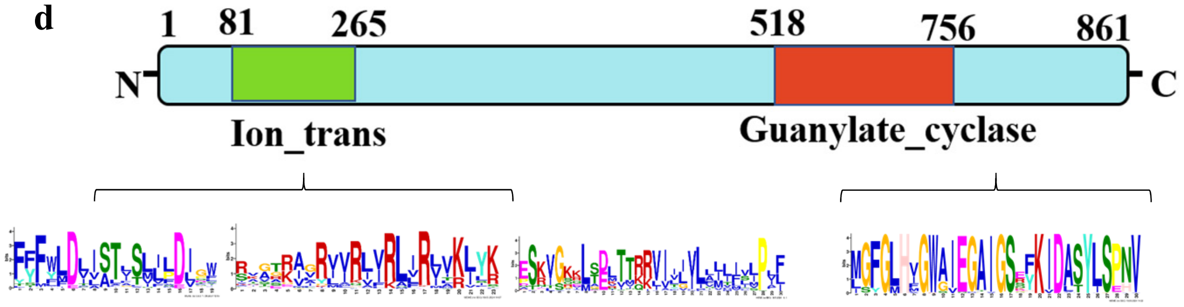


**Supplementary Figure 1**
