## Supplementary Data 1 for "Membrane Proteome of *Phytophthora parasitica* Zoospores: How Does Sensing Occur?"

**Liquid chromatography and mass spectrometry analysis**

M-F and M-CB samples were fractionated by electrophoresis under reducing conditions into one-dimensional Sodium Dodecyl Sulfate-polyacrylamide (10%) gels. Four biological replicates of each fraction were loaded into a gel (5 µg/µL), subjected to electrophoresis (100V, 60 min) and later stained with Coomassie blue (Thermo Scientific PageBlue Protein Staining Solution). Gel tracks corresponding to biological replicates were sectioned into several bands. Each band was then cut into 1 $mm^3$ cubes and subjected to a washing process using water and acetonitrile (ACN). Subsequently, the gel pieces were reduced with 10 mM dithiothreitol in 50 mM ammonium bicarbonate for 30 min at 56°C. Following reduction, the samples were alkylated with 55 mM iodoacetamide in 50 mM $NH_4HCO_3$ for 20 min at room temperature in the dark.

The gel cubes were washed with 50 mM $NH_4HCO_3$ for 10 min, followed by a 15-min wash with acetonitrile (ACN), and then dried for 2 min in a SpeedVac (Savant, Apeldoorn, Netherlands). The cubes were subsequently incubated overnight at 37°C in a solution containing 25 mM $NH_4HCO_3$, 5 mM $CaCl_2$, and 12.5 ng/µL sequencing-grade modified trypsin (V5111; Promega, Madison, WI, USA)[18]. Enzymatic digestion was terminated by adding 5% formic acid (FA). The resulting peptides were extracted by washing the cubes twice with ACN. The samples were then dried for approximately 2 hours and desalted using OMIX C18 pipette tips (100 µL, A57003100) (Agilent, Santa Clara, CA, USA).

Samples were analyzed using a nanoUHPLC system (nanoElute) coupled to a TimsTOFpro mass spectrometer (Bruker Daltonics, Germany). Ten microliters of each sample were injected and separated on a reverse-phase C18 column with an integrated CaptiveSpray Emitter (75 µm ID x 250 mm, 1.6 µm, Aurora Series with CSI, ionOpticks, Australia) at a flow rate of 200 nL/min. The mobile phase consisted of 0.1% FA in water (Phase A) and 0.1% FA in ACN (Phase B). The liquid chromatography gradient started at 2% Phase B, ramping to 5% B in 1 min, then to 13% B in 18 min, followed by 22% B in 11 min, and finally to 95% B in 3 min, held for 7 min.

The TimsTOFpro mass spectrometer was operated with the CaptiveSpray nano-electrospray ion source. The source temperature was set to 180°C, with a spray voltage of 4500 V. Nebulizer gas (nitrogen) was supplied at 0.4 bar, and dry gas (nitrogen) was set at a flow rate of 3.0 L/min. MS and MS/MS data were acquired in positive polarity using Parallel Accumulation-Serial Fragmentation (PASEF)[19] Data Dependent Acquisition (DDA) mode, with 10 PASEF MS/MS scans per cycle and a 100% duty cycle (Parallel Accumulation TIMS, EP3054473A1) (Parallel Accumulation TIMS, US9683964B2)[20]. Peptides were detected over a mass range of 100 to 1700 m/z, with a target intensity of 20,000 and an intensity threshold set at 2500. The collision energy was ramped linearly as a function of mobility, from 59 eV at 1/K0 = 1.3 Vs/cm² to 20 eV at 1/K0 = 0.7 Vs/cm² (TimsControl version 2.0.53.0).

The acquired DDA spectra were initially examined using Data Analysis software (version 5.3, Bruker Daltonics, Germany). Subsequently, the data were processed with PEAKS Studio (version Xpro, Bioinformatics Solutions) against the proteome predicted from the *P. parasitica* 310 genome[21]. The PEAKS identification search allowed for one missed cleavage, with carbamidomethylation set as a fixed modification and methionine oxidation as a variable modification. Contaminants were removed, with a parent mass error tolerance set to 20 ppm and a fragment mass error tolerance of 0.02 Da. Only proteins identified with an FDR of 1%, with at least one unique peptide, were selected.

Relative quantitative analysis of proteins was performed using the PEAKS Q label-free quantification method in PEAKS. This method, similar to the approach described by MaxQuant for high peptide identification rates and proteome-wide protein quantification[22], was applied to all four biological replicates for both groups M-F and M-CB. According to the method indications, for quantification results, only proteins with an FDR of less than 1% and a fold change greater than 2 were selected. Peptides considered for quantification had to be identified in both groups and detected in at least two samples per group, with quantification requiring a minimum of one peptide.
